# Foot scales in the Early Cretaceous bird *Gansus yumenensis* from China

**DOI:** 10.1101/2021.06.07.447457

**Authors:** Tao Zhao, Zhiheng Li, He Zhang, Yanhong Pan

## Abstract

Most modern birds have scales covering the foot and feathers elsewhere. Discoveries of fossil feathers attached to the metatarsus in non-avian dinosaurs and basal birds suggests that the avian scales are secondarily derived from feathers. However, our knowledge of early avian scales and their taphonomy is still limited, due to the scarcity of fossil record. Here we employ multiple techniques to characterize the morphological and chemical details preserved and investigate how they are preserved in the skin of IVPP V15077, a referred specimen of the Early Cretaceous *Gansus yumenensis*. Results show that two types of scales, scutellate and interstitial scales, are preserved in IVPP V15077, which, in combination with previous discovery of scutate and reticulate scales in other Early Cretaceous birds, indicates that all four types of scales present in modern birds have appeared in the Early Cretaceous. SEM observations and Raman analysis suggest that the skin of *Gansus yumenensis* may be pigmented. Elemental mapping indicates that aluminosilicates and calcium phosphate are involved in the mineralization of the skin.

## Introduction

Most modern birds have scales covering the foot (tarsometatarsus and toes) and feathers covering most of the rest of the body, with several wild species and some breeds of domestic pigeon and chicken display foot feathering (ptilopody) [1–3]. The scales can be categorized into four types: scutate scales that are large and somewhat overlapping on the anterior surface of the tarsometatarsus and dorsal surface of toes, scutellate scales that are somewhat smaller than scutate scales and located on the caudal surface of the tarsometatarsus, reticulate scales that are located on the plantar surface of toes, and interstitial scales that are morphologically similar to reticulate scales but are located on the tarsometatarsus [4]. The scutate, scutellate, and interstitial scales have similar patterns of keratinization, containing both α-keratins and β-keratins; the reticulate scales, by contrast, contain α-keratins but no β-keratins [4]. The interstitial scales are also referred to as reticulate scales in some studies [e.g., 5–7].

Fossil discoveries of non-avian dinosaurs and basal birds with feathers attached to the metatarsi suggest that foot feathering is the primitive state for birds [8–11], which is consistent with the view that avian scales are secondarily derived from feathers [12–14]. The foot feathering in modern birds might represent a reversion to the ancestral state [15]. Another view concerning the origin of avian scales is that they are homologues of reptilian scales [5,16]. To date, our knowledge of the foot scales in early birds is limited, due to the scarcity of fossil record [11,17–19].

IVPP V15077 (Institute of Vertebrate Paleontology and Paleoanthropology) is a referred specimen of *Gansus yumenensis* with scales well preserved *in situ* around the joint of the tibiotarsus and the tarsometatarsus [17]. To date, all reported specimens of *Gansus yumenensis* are from Xiagou Formation, Changma basin, China [17,20,21]. Stable isotope chemostratigraphy places the age of the bird quarries in the early Aptian [22]. Despite the well preservation of the skin in IVPP V15077, no investigation on the ultrastructure and the chemistry of the skin has been performed. The aim of the present study is to investigate what morphological and chemical details are preserved in the skin and how they are preserved by employing multiple techniques, including scanning electron microscopy (SEM), scanning electron microscopy-energy dispersive X-ray spectrometry (SEM-EDS), Raman spectroscopy, and X-ray powder diffraction (XRPD).

## Material and Methods

One skin sample was removed from IVPP V15077 using a sterile blade and directly analyzed using scanning electron microscopy (SEM), scanning electron microscopy-energy dispersive X-ray spectrometry energy dispersive X-ray spectrometry (SEM-EDS), and Raman spectroscopy. One sediment sample was scraped from the bedding plane surface where the fossil is preserved, and powered for X-ray powder diffraction (XRPD) analysis.

SEM observations were performed using a Sigma 500 Field Emission Scanning Electron Microscope (FE-SEM) at 1.5 keV. Elemental mappings of the skin sample were performed using a Tescan MAIA3 FE-SEM equipped with an Energy Dispersive X-ray Spectrometry (EDS) at 8 keV and at 20 keV.

Raman analysis of the skin sample was performed using a LabRAM HR Evolution Raman spectrometer with a 532.11 nm laser and a 50 × Olympus objective with a long working distance. A 600 groove/mm grating was used with the spectral resolution better than 2 cm^−1^. Spectra were acquired with four accumulations and accumulation time of 4s to 8s. Raman analysis was also performed a black chicken feather to obtain spectra of eumelanin [23] for comparison.

XRPD analysis of the sediment sample was performed using a Bruker D8 ADVANCE diffractometer with Cu Kα radiation and the scanning angel ranged from 3° to 90° of 2θ.

## Results

The skin preserved around the joint of the right tibiotarsus and tarsometatarsus is exposed in the inner view (Figure 1). The scales are non-overlapping. The proximal scales are large and elongate, and correspond to the scutellate scales. The distal and medial scales are small and rounder than the proximal scales, and correspond to the interstitial scales. The regions between scales, referred to as sulci in extant birds, are lighter in color than the scales, as seen from where the sulci are exposed (Figure 2A). Most part of the sulci is covered by a layer of sediment.

**Figure 1.**
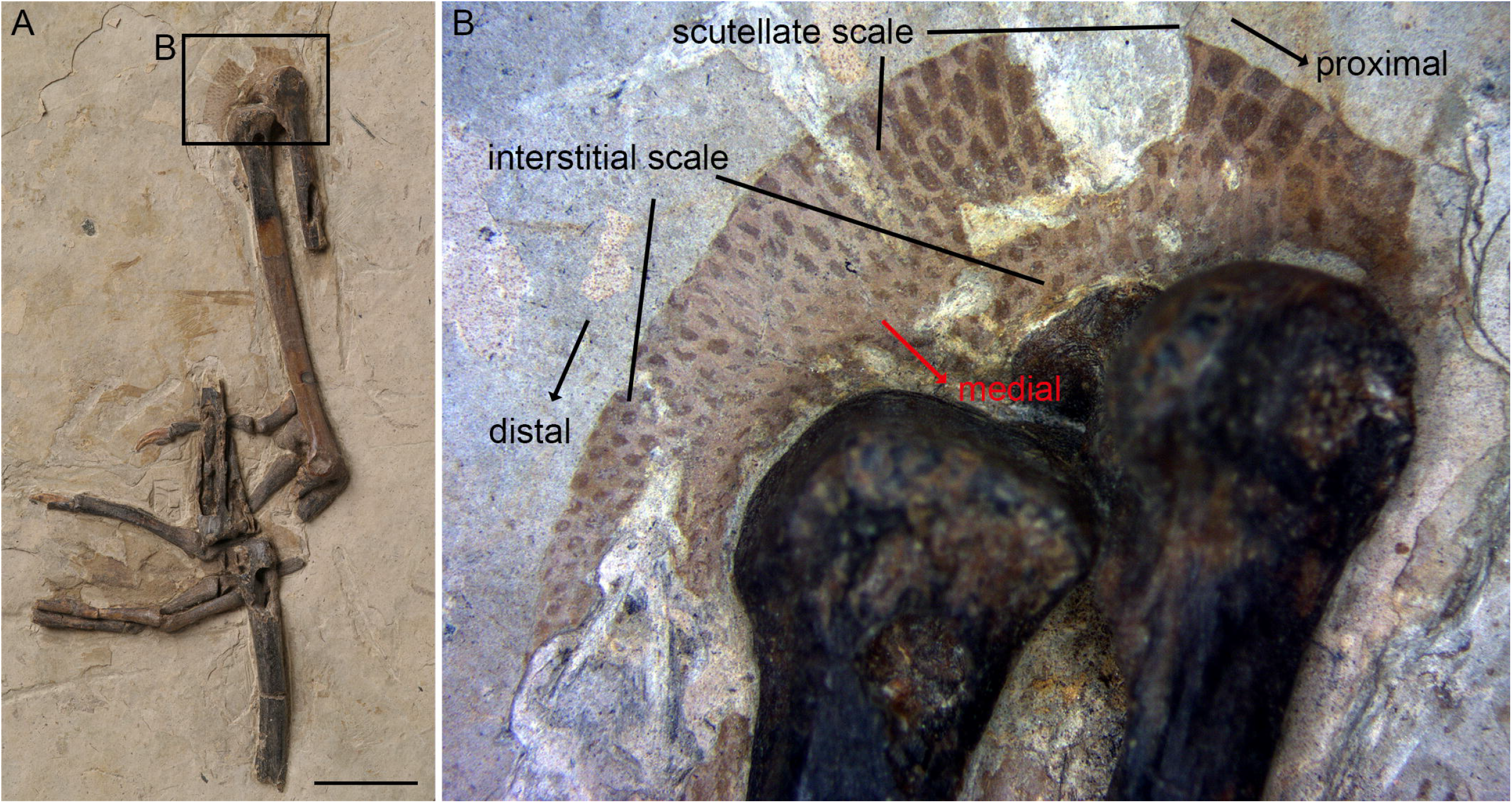
Photo of IVPP V15077, a referred specimen of *Gansus yumenensis* showing the skin preserved around the joint of the right tibiotarsus and tarsometatarsus. Scale bar in (A) equals to 1 cm.

**Figure 2.**
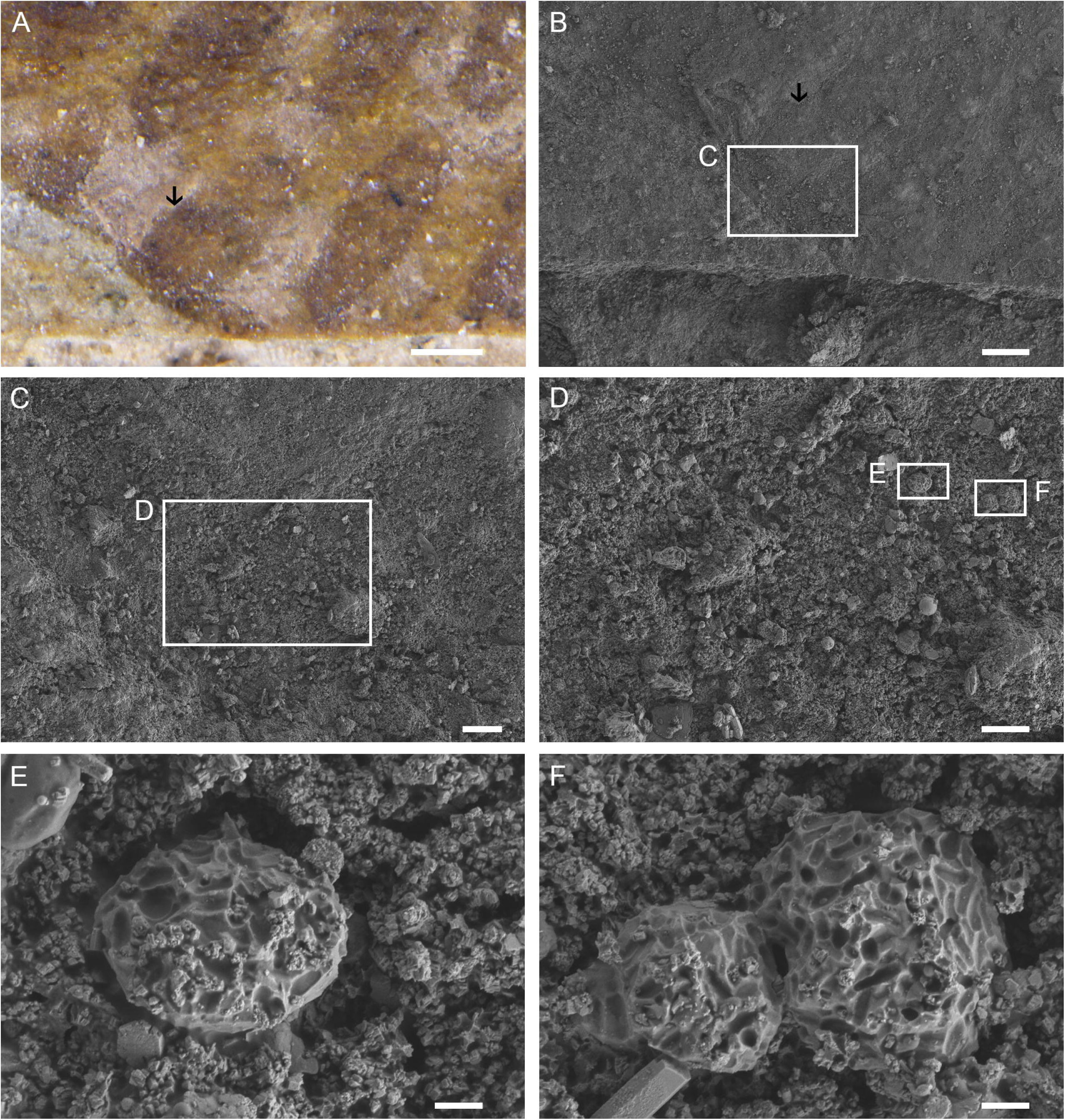
Photo and SEM images of the skin preserved in IVPP V15077. Arrows in (A) and (B) mark the same position. Scale bar equals to 200 μm in (A), 100 μm in (B), 20 μm in (C), 10 μm in (D), 1 μm in (E) and (F).

**Figure 3.**
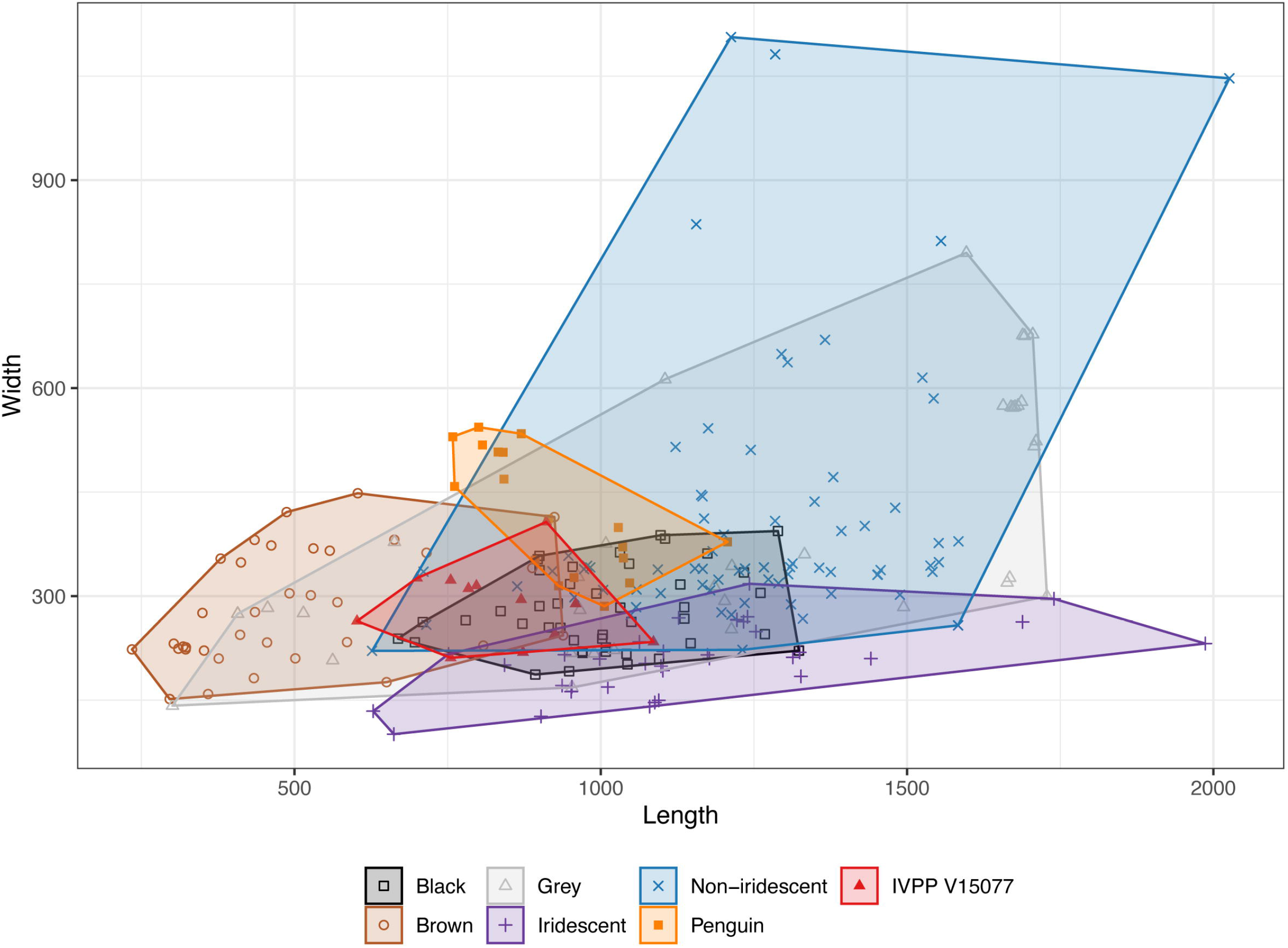
Comparison of the length and width of the microbodies in the skin of IVPP V15077 with those of melanosomes from feathers. Data on melanosomes from feathers are from ref. 31. Feathers are categorized into 6 color groups: black, brown, grey, iridescent, non-iridescent structural color and penguin.

SEM reveals impressions of ovoid and rod-like microbodies on some scattered fragments in the skin (Figure 2). Such microbodies likely represent melanosomes following recent studies on fossil feathers and reptile skin [24–30]. The dimensions of these microbodies fall within the range of melanosomes from feathers, more specifically, the overlapping region of melanosomes from black feathers, gray feathers, brown feathers, and feathers with non-iridescent structural colors [31].

The elemental mapping was performed with two accelerating voltages of 8 keV and 20 keV to detect the vertical variation of the elemental distribution. The elemental mapping at 8 keV show that the scales have an elevated concentration of C than the sulci. This discrepancy is less distinct in the elemental mapping at 20 keV, indicating the C is distributed near the surface. The layer of sediment covering the sulci has an elevated concentration of Ca and P, indicating the presence of calcium phosphate. Where this layer of sediment was removed (arrows in Figure 4), the elements Si, Al, Fe, Na, S, Mg, and K show no spatial partitioning between scales and sulci.

**Figure 4.**
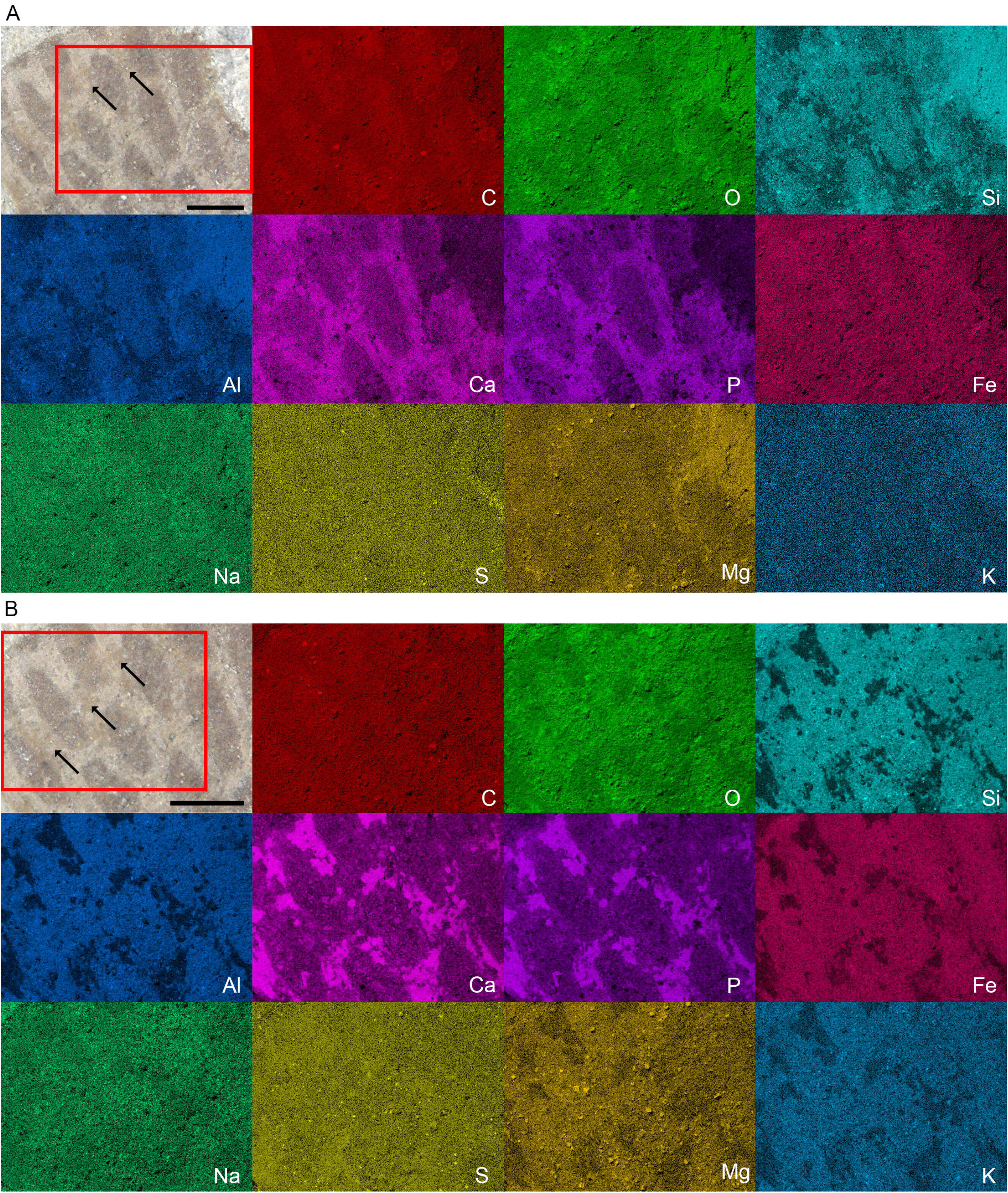
Elemental maps for the skin in IVPP V15077 at 8 keV (A) and 20 keV (B). Arrows indicate where the layer of sediment covering the sulci was removed. Scale bar equals to 500 μm in (A) and (B).

**Figure 5.**
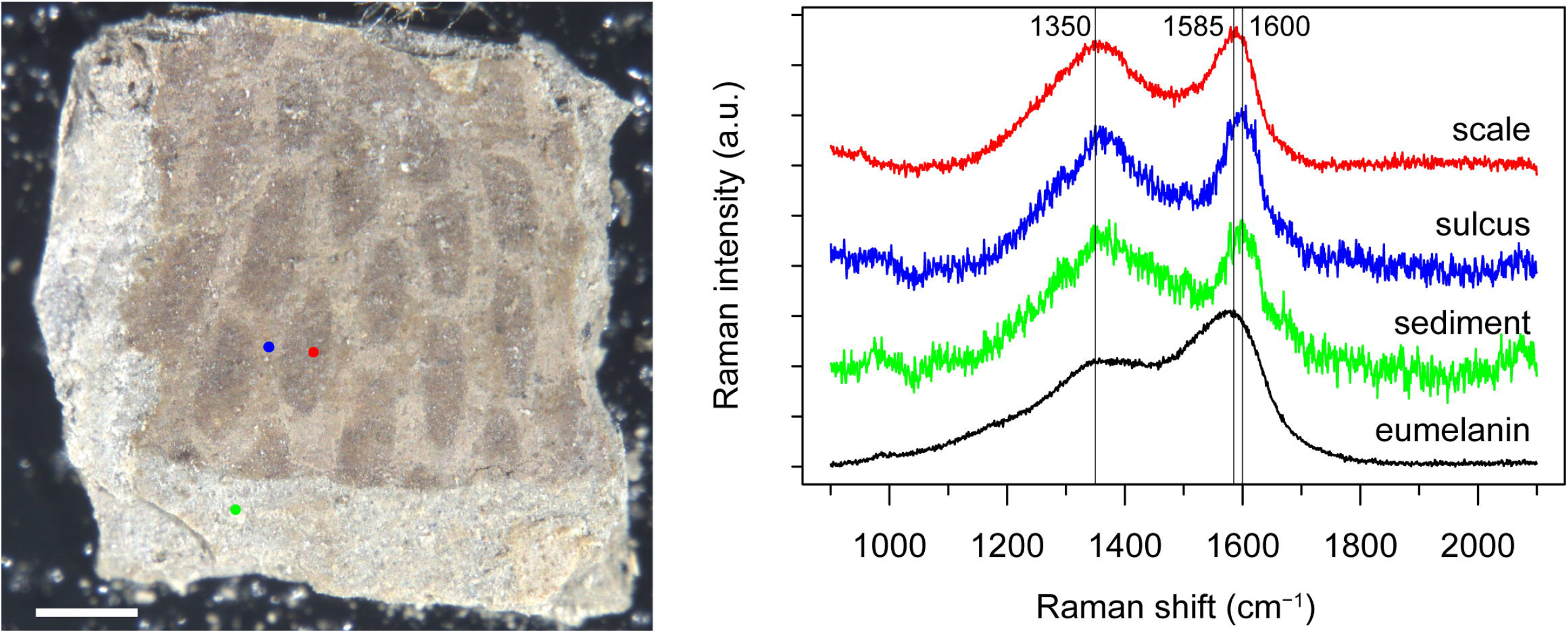
Raman spectra of the skin in IVPP V15077. Scale bar equals to 500 μm.

Raman spectroscopy confirms the presence of carbonaceous material in the fossil skin. The Raman spectra of the sulcus and the sediment are similar; both exhibit two broad bands near1350 cm^−1^ and 1600 cm^−1^, which are characteristic of carbonaceous materials and referred to as the D-band and the G-band. By contrast, the G-band in the Raman spectrum of the scale has a lower wavenumber (1585 cm^−1^), resembling that in eumelanin.

XRPD shows that the sediment consists mainly of analcime, apatite, quartz, calcite, and illite (Figure 6).

**Figure 6.**
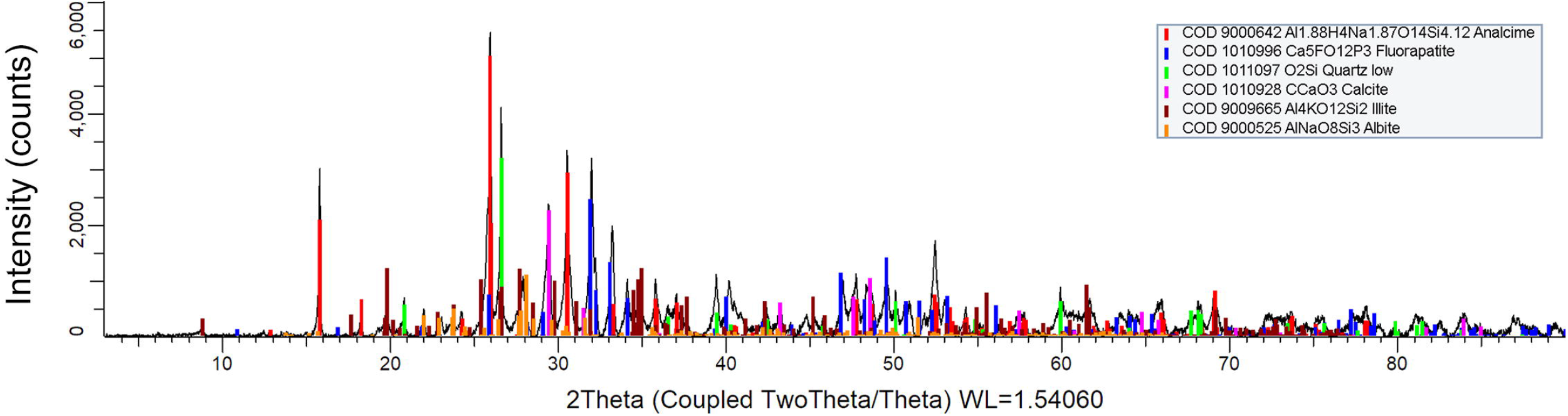
X-ray powder diffraction analysis of the sediment from the bedding plane surface where IVPP V15077 is preserved.

## Discussion

### Foot scales in early birds

Fossil record is key to determining when modern taxa acquire their morphological characters. Among the four types of scales in modern birds, reticulate scales on the plantar surface of toes appeared earliest in the lineage leading to modern birds, and have been found in basal birds *Sapeornis* [32] and *Confuciusornis* [18]. Discoveries of metatarsal feathers in basal birds *Sapeornis* and *Confuciusornis* and non-avian dinosaurs imply that scales on the tarsometatarsus in modern birds evolved secondarily from feathers in the Ornithuromorpha [11], the most inclusive avian clade that contains all modern birds but not Enantiornithes [33]. In the basal ornithuromorph *Yanornis* from the Early Cretaceous Jehol Biota, scutate scales covering the anterior surface of the tarsometatarsus and dorsal surface of toes have been previously found [11]. Reexamination of IVPP V15077 here reveals the presence of scutellate scales and interstitial scales in the Early Cretaceous ornithuromorph *Gansus yumenensis*. These results show that all the four types of scales in modern birds [4] have appeared in the Early Cretaceous.

### Pigmentation of the preserved skin

SEM observations and Raman spectroscopy suggest that the preserved skin of *Gansus yumenensis* is likely pigmented. The dimensions of the melanosome-like structures fall within the range of melanosomes from feathers [31]. It has been noted that the Raman spectra of eumelanin resemble that of disordered graphite [23,34]. Raman spectroscopy was thus cautioned to be used alone as a definitive test for the presence of eumelanin in fossils [35], as organic matter can be progressively transformed into graphite when heated [36]. Nevertheless, the assignment of the 1585 cm^−1^ band in the Raman spectrum of the scale to eumelanin here is supported by its wavenumber difference from the G-band in the spectrum of the sulcus. Raman spectral parameters of organic matter is indicative of the temperature experienced by host rocks and the thermal maturity of organic matter [37,38]. A G-band at 1585 cm^−1^ indicates a higher maturity level than a G-band at 1600 cm^−1^ band. As the scales and the sulci have the same geological history, their difference in Raman spectra supports an original compositional difference.

### Taphonomy

Usually, the skin decays quickly after the death of birds [39]. The preserved carbonaceous matter in the fossil suggests that the skin was stabilized from enzymatic and other postmortem changes soon after the death of the bird [40].

Sedimentary facies analysis indicated that the Xiagou formation in Changma Basin in dominated by lacustrine deposits [41]. The primary to early diagenetic dolomites in this Formation suggests that the lake is closed, alkaline to saline [42]. The identification of analcime in the sediment is consistent with this interpretation of the lake. Reaction of alkaline lake water and detrital clay and perhaps other materials can result in the formation of analcime [43].

The pattern of elemental distribution suggests two mineralization modes are responsible for the preservation of the skin in IVPP V15077. The epidermis, including the scales and sulci, is mineralized in aluminosilicates. As the skin of IVPP V15077 is exposed in the inner view, the layer of carbonate phosphate covering the sulci is supposed to correspond to the dermis. Preservation of soft tissues, including skin, in calcium phosphate is relatively common in fossil record [44,45]. To date, research into mineralization of soft tissues in aluminosilicates is mainly focused on marine settings, aiming to understanding the Burgess Shale-type fossilization [e.g., 46–49]. Al is suggested to be a key factor in the stabilization of soft tissues [47,48,50]. It has been hypothesized that Al can induce taphonomic tanning, a concept borrowed from leather industry, involving secondary cross-linking of structural biomolecules that protects them from bacterial degradation [47]. The surrounding sediment can serve as the source of Al. Experimental work demonstrated that the decay of soft tissues can lower the pH values and lead to the acid hydrolysis of clay minerals in the surrounding environments both in marine settings [48] and in fresh water settings [50]. The experimental work simulating marine environments demonstrated that kaolinite and chlorite sediment enhanced soft tissue preservation compared with sediment-free control and the effect of kaolinite is higher than that of chlorite [48]. Consistent with the hypothesis of taphonomic tanning of Al is that more Al was released during the acidic hydrolysis of kaolinite compared to chlorite [48]. Likewise, the experimental work simulating fresh water environments demonstrated that kaolinite and montmorillonite sediment enhanced soft tissue preservation compared with sediment-free control and the effect of kaolinite is higher than that of montmorillonite [50]. Though recent studies focused on the role of kaolinite in the preservation of soft tissues [48–50], kaolinite is by no means the only mineral that can produce Al through acid hydrolysis. Both albite and analcime identified in the sediment from the bedding plane of IVPP V15077 can be dissolved by organic acid [e.g., 51,52]. Analogously, Fe-based tanning may also contribute to the inhibition of the decay of soft tissues [47,53].

## Acknowledgements

We thank Yaxiao Wang (Key Laboratory of Surficial Geochemistry, Ministry of Education, School of Earth Sciences and Engineering, Nanjing University) for assistance during SEM observations and Raman analysis, Yan Fan (State Key Laboratory of Palaeobiology and Stratigraphy, Nanjing Institute of Geology and Palaeontology, Chinese Academy of Sciences) during SEM-EDS analysis, and Yuguan Pan (State Key Laboratory for Mineral Deposits Research, School of Earth Sciences and Engineering, Nanjing University) during XPRD analysis.

